# Spatial replication can best advance our understanding of population responses to climate

**DOI:** 10.1101/2022.06.24.497542

**Authors:** Aldo Compagnoni, Sanne Evers, Tiffany Knight

## Abstract

Understanding the responses of plant populations dynamics to climatic variability is frustrated by the need for long-term datasets. Here, we advocate for new studies that estimate the effects of climate by sampling replicate populations in locations with similar climate. We first use data analysis on spatial locations in the conterminous USA to assess how far apart spatial replicates should be from each other to minimize temporal correlations in climate. We find that on average spatial locations separated by 316 Km (SD = 149Km) have moderate (0.5) correlations in annual precipitation. Second, we use simulations to demonstrate that spatial replication can lead to substantial gains in the range of climates sampled during a given set of years so long as the climate correlations between the populations are at low to moderate levels. Third, we use simulations to quantify how many spatial replicates would be necessary to achieve the same statistical power of a single-population, long-term data set under different strengths and directions of spatial correlations in climate between spatial replicates. Our results indicate that spatial replication is an untapped opportunity to study the effects of climate on demography and to rapidly fill important knowledge gaps in the field of population ecology.

## INTRODUCTION

Understanding the responses of biodiversity to climate drivers is necessary to mitigate and adapt to climate change (Urban et al. 2016). In recent years, the field of ecological forecasting has experienced substantial theoretical and applied growth (Lewis et al. 2022). These forecasts on population dynamics are necessary to assess species extinction risk (Mace et al. 2008) and predict range shifts (Schurr et al. 2012). However, forecasting the effects of climate on populations remains elusive. This plausibly occurs because it takes 20-25 years of data to sufficiently describe the relationship between climate and demography (Teller et al. 2016, Tenhumberg et al. 2018). This large replication is necessary to sample a sufficiently wide range of climatic conditions and to increase statistical power.

We have important knowledge gaps in population ecology that cannot wait 20-25 years to be filled. Our recent synthesis showed that knowledge on climate-demography relationships for plants is particularly poor for the species-rich tropics, and for species with extreme generation times (Compagnoni et al. 2021b). We need immediate research targeting these locations and plant life histories. As most plant ecologists are at young career stages, we need to engage Doctoral Researchers and Postdoctoral Associates in this field of research. However, this will not happen in case projects last decades.

We propose that new studies should prioritize spatial over temporal replication to assess the demographic responses of a species to climate. Having both spatial and temporal data allows collecting a high sample size in a relatively short period of time. Spatial sampling increases our statistical power by increasing the range of climates sampled per unit time, and allowing us to “see through the noise” caused by non-climatic factors. The range of climates sampled increases as the distance between populations decreases the correlation of yearly climatic anomalies among them.

We are not advocating a “space-for-time substitution”, but to estimate the short-term effects of climate by prioritizing spatial versus temporal replication. Space-for-time substitutions infer the long-term effects of climate on plant populations (Blois et al. 2013). These long-term effects are typically larger than short-term climate effects, possibly because they include the effects of indirect effects (e.g. on biophysical conditions, Elmendorf et al., 2012). Instead, short-term climatic effects are directly relevant to inform the management and conservation of populations during the upcoming century of rapid climate change (e.g. Compagnoni, Pardini, et al., 2021; Hunter et al., 2010). Here, we therefore advocate to replicate sampling across populations that occur in similar climates, and to use this spatial data as *replicates* of the same temporal process. This recommendation relies on the assumption that in similar environments, plant populations should respond similarly to climate anomalies. Such assumption has so far motivated much research on the demography across species ranges (Kleinhesselink & Adler, 2018; Morley et al., 2017). However, to our knowledge, no study on climate-demography relationships has yet prioritized spatial replication across sites with similar climatic conditions.

In this manuscript, we show the opportunities of spatial replication in climate-demography studies through data analysis and simulation. First, we assess how far apart populations must be from each other to attenuate temporal correlations in climate using gridded climatic data from the conterminous USA. Second, we consider how sampling design can maximize the range of climates captured during a study. To do so, we estimate how the range of climates sampled depends on the climate correlations among the populations and the study duration (between five and 30 years).

Third, we use simulation to quantify the statistical power of a climate-demography relationship across sampling designs that vary in the number of spatial and temporal replicates, and in the strength of spatial correlations in climate between populations. In this simulation, we also address cases in which populations respond differently to climate. Based on these results, we make recommendations for new demographic data collection efforts.

## METHODS

### Spatial correlation in climate

To understand how spatial correlation in climate depends on the distance between sites, we estimate the spatial correlation of annual climate in the conterminous United States of America (USA), a large and climatically heterogeneous section of terrestrial environments. We downloaded monthly temperature and precipitation data for the conterminous USA from the CHELSA database (Karger et al. 2017, 2018). CHELSA data is accurate on varied topographic terrain, and gridded climatic data generally correlates strongly with weather station data (Behnke et al. 2016). We downloaded data between 1979 and 2013 following a regular grid of 0.5 degrees, for a total of 3261 locations. For each location, we calculated annual temperature means, annual precipitation sums, and computed their standardized yearly anomalies (z-scores, henceforth “anomalies”).

For each location, we calculated the average distance at which the correlation in yearly climate anomalies is expected to decrease to 0.5 or lower. To do so, we calculated the correlation between the 35 annual anomalies at a location, and the same anomalies observed at the remaining 3260 locations. Then, we fit a local polynomial regression using observed correlations as a response variable, and distance from the original location as predictor. The predictions of this local regression identified the distance at which the correlation in climate anomalies is expected to be 0.5 or lower. Finally, we produced heatmaps showing, for each location, the average predicted distance at which the correlation in climatic anomalies is 0.5 or lower.

To show a sample of the raw data we used for this analysis, we show the data referred to five reference locations. We picked these locations subjectively, locating them in the Southwestern coast, Northwestern coast, Northeastern Coast, Southeastern coast, and in the center of the USA. For each location, we first plotted the correlation between climatic anomalies versus the distance from the reference location, including the average prediction of the polynomial regression summarizing this relationship. Then, we produced heatmaps showing how the climatic anomalies observed across the USA correlated with the reference location.

### Range of climates sampled with different sampling designs

We performed simulations to understand how the range of climate anomalies sampled changes depending on temporal replication, spatial replication, and the spatial correlation of climatic anomalies. We estimated the range of climatic anomalies sampled at two sites using a multivariate normal (MVN) distribution:

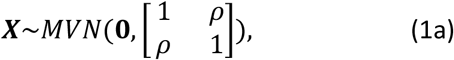

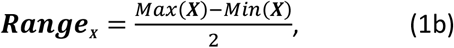

where **X** is an *n* by 2 matrix of climate anomalies at two separate sites, *n* is the temporal replication of the study, *ρ* represents the correlation between the climatic anomalies of the two sites. The parameters of the MVN distribution show **0** for the means, and diagonal elements of 1 for the variance to simulate a series of anomalies. When simulating the climate at a single site, we needed a single series of climate anomalies, so we substituted Eq. (1a) with *X*∼*Normal*(0,1). To obtain ***Range***_***X***_, the expected range of **X** values (Eq. 1b), we simulated Eq. 1a 1000 times across *n* values ranging from two to 30 in increments of one, and across a series of *ρ* values of 0, 0.5, 0.95, and one. Finally, we calculated the mean of **Range**_**X**_ across these 1000 replicate simulations.

### Statistical power for climate-demography relationship with different sampling designs

We used simulations to quantify the statistical power of the relationship between climate and population growth rate for different spatio-temporal sampling designs and different spatial correlations in climate. Starting from a case with two separate sites,

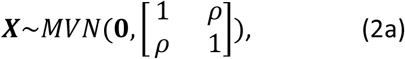

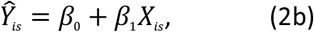

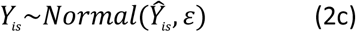

where ***X*** is an *n* by 2 matrix of normally distributed climatic anomalies, *n* refers to temporal replicates, *Ŷ*_*is*_ is the average prediction of the model referred to site *s* and year *i, β*_0_ the intercept of the linear model, *β*_1_ is the slope, and *Y*_*is*_ represents the observed log population growth rate. *Y*_*i*_ is a log population growth rate because we have synthetic estimates of climatic effects on this variable (Compagnoni et al. 2021b), because it is the central focus of demographic theory (Sibly and Hone 2002), and because it is normally distributed, facilitating simulations and their interpretation. For this and subsequent simulations we used a *β*_1_ value of 0.05, and a *ε* value of 0.15 to reflect empirical estimates from 162 plant populations (Compagnoni et al. 2021b). We simulated the process described in Eq. 2 1000 times using study durations, *n*, of three, four, and five years, correlations *ρ* of 0.95 and 0.5, 0, and -0.5 and a number of spatial replicates of two, 10, 20, 30, 40, and 50. We divided spatial replicates in two subsets, and assigned each subset to one of the two series of climatic anomalies simulated by Eq. 2. For example, when spatial replicates were 50, we subdivided these replicates in two groups of 25 replicates, each group experiencing identical climate. We used low values for *n* to reflect that the median length of demographic studies of plants is four years (Salguero-Gómez et al. 2015). This sampling effort likely reflects the length of many PhD programs.

As an alternative, we also simulated assuming a single censused population, using study durations of 20 and 30 years. This simulation was meant to represent the handful of single site, long-term studies found in the literature (e.g. Chu et al. 2016). In these simulations, we substituted Eq. 2a with *X*∼*Normal*(0,1). We calculated power as the proportion of the 1000 simulations for which *β*_1_ had a p-value below 0.05. This power analysis based on simulation is a straightforward way to quantify how the uncertainty of model estimates is influenced by the sampling design.

The power estimate described above assumed that *β*_1_, in Eq. 2b, were the same for each population. Therefore, we also addressed the sensitivity of our power estimates to spatial variation in *β*_1_. We ran all of our analyses in R version 4.0.2 (R Core Team 2020). We fit polynomial regressions using the R function loess, and we produced spatial plots using package ggmap (Kahle and Wickham 2013). We ran the analyses on the spatial correlation in climate using a high-performance computing cluster.

## RESULTS

### Spatial correlation in climate

In the conterminous USA annual temperature anomalies are strongly correlated even at relatively large (e.g. 500Km, Fig. S1-5A) distances. The correlation between precipitation anomalies is less strong, and it decays more rapidly with distance (Fig. S1-5C). In the conterminous USA, our loess models predict that the distance to reach a correlation of 0.5 is on average 316 Km (SD = 149 Km) for precipitation and 1460Km (SD = 428Km) for temperature. However, there is substantial heterogeneity in the decay of correlation for each location (Fig. 1). Considering the 5^th^ and 95^th^ percentiles of the distances at which the correlation of climate anomalies is predicted to be 0.5, these range from 52 to 568 Km for precipitation, and from 868 to 2196Km for temperature. Sites with a predicted correlation of zero are on average always at least 1000Km apart: only 5% of the predicted distances at which the correlation is zero have a distance equal to or less than 1000 and 1920 Km for precipitation and temperature, respectively.

**Figure 1.**
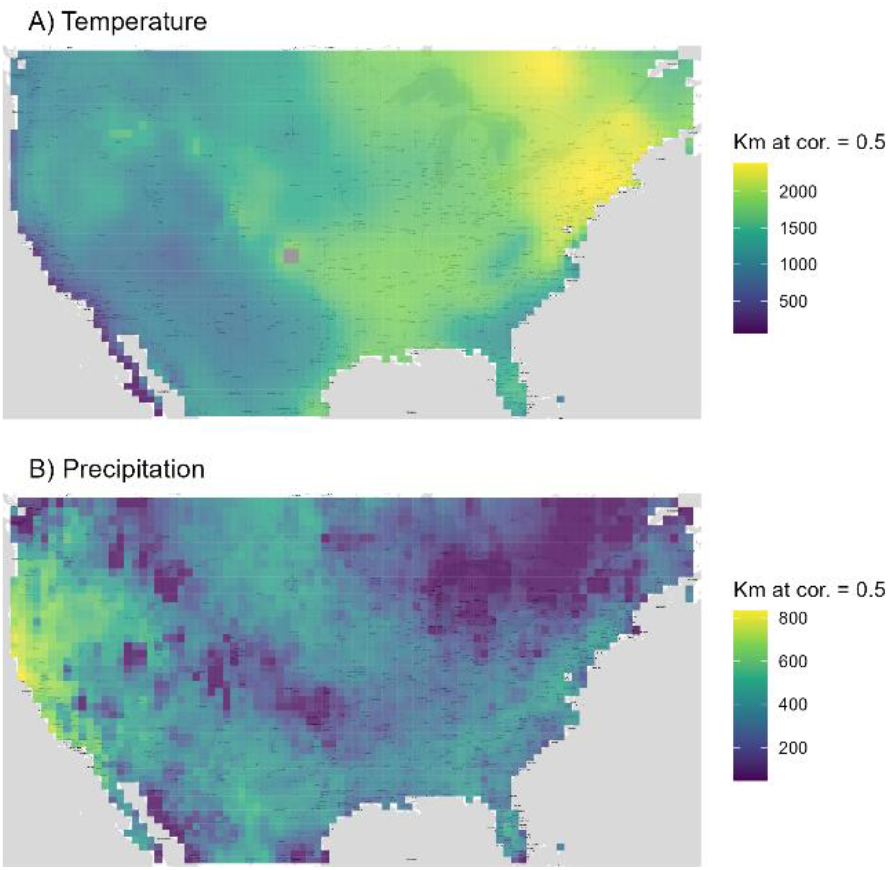
The correlation between temperature and precipitation anomalies decays slowly with distance. The heatmaps show for each grid cell, the average distance at which the correlation in temperature (A) and precipitation (C) anomalies decreases to 0.5 or lower. These average distances are estimated using a local regression (loess) model.

### Range of climates sampled with different sampling designs

Spatial replication can lead to substantial gains in the range of climates sampled during a given set of years so long as the climate correlations between populations are at low to intermediate levels. To reach a range of ±2 standard deviation at a single site, one would on average need 27 years of data. This number of years decreases to 20 when using two sites whose climate has correlation 0.9, and 15 when two sites have correlations 0.5 (Fig. S6).

### Statistical power for climate-demography relationship with different sampling designs

Our power analysis indicates that spatial replication greatly increases the power to detect a relationship between climate and population growth rate (Fig. 2). The statistical power of very long time series for a single site is comparable to that of datasets with high spatial replication. One site sampled for 20 and 25 times provides a statistical power of about 30 and 41%, respectively. These two statistical powers are reached with just three temporal samples distributed across, respectively, 10 and 20 spatial replicates experiencing medium to highly correlated climate (0.5, 0.95). With four temporal samples and 10 spatial replicates, a statistical power of 40% is reached with climate correlation 0.95; with five temporal samples and 10 spatial replicates, statistical power always exceeds 40%. Lowering climatic correlation between populations from 0.95 to 0.5 slightly increases statistical power: depending on the number of temporal samples and spatial replicates, increases in statistical power vary from 4 to 8% (Fig. 2).

**Figure 2.**
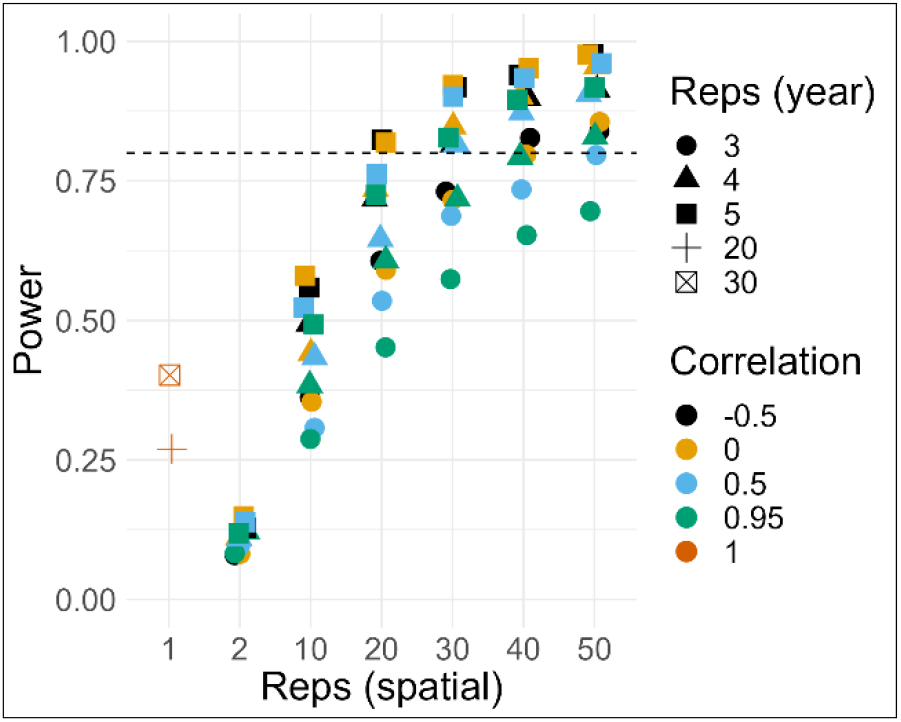
Spatial replication provides a statistical power similar, or higher, to temporal replication. Plot showing statistical power on the y-axis against spatial replication on the x-axis. Symbols show temporal replications of three (circles), four (triangles), five (squares), 20 (cross), and 30 (crossed square). The color of symbols refers to the correlation among spatial replicates. This correlation is one for the simulations with a single spatial replicate. The dashed horizontal line denotes a statistical power of 80%.

Variation in the effect of climate on demography (represented by *β*_1_ values) did not noticeably affect statistical power (Fig. S7). Statistical power remained unchanged, reflecting that is was influenced by the average *β*_1_, which was still 0.05, rather than its variation.

## DISCUSSION

Until recently, understanding the response of plant species to climatic variation has relied on either long-term monitoring efforts, which are rare (Salguero-Gómez et al. 2015), or on responses of plant populations to spatial climate gradients (“space-for-time substitutions”, Blois et al., 2013) which, are affected by several confounding factors. Specifically, spatial climatic gradients do not measure the direct effect of climate on populations, because they encompass substantial changes in community composition (Whittaker 1970), and biophysical factors (Shaver et al. 2000). Fortunately, our power analysis shows that we can propel our understanding of species responses to climate using spatial replicates that come from sites with similar long-term average climates.

We argue that using numerous spatial replicates in studies of demographic responses to climate is an opportunity that should be exploited urgently by conservation scientists. Our power analysis suggests that large spatial replication allows obtaining estimates of climate effects in as little as three years. This short temporal horizon would allow early career researchers or distributed networks to study the effects of climatic variability on populations. Such abundance of investigators would facilitate ameliorating the taxonomic, geographic, and life-history biases present in current data (e.g. Compagnoni et al., 2021). The result would encourage the development and improve the quality of climate change vulnerability assessments. The IUCN recommends evaluating vulnerability by focusing on time series of population data (Mace et al. 2008). These data can be readily used to estimate the effects of climate on population dynamics and, as a result, climate vulnerability (Hunter et al. 2010, Jenouvrier et al. 2022). However, given that long-term population data are rare in plants (Salguero-Gómez et al. 2015), large spatial replication provides an opportunity to perform such population-based climate vulnerability assessments.

We suggest new studies focusing on sites with similar climate located at the leading or rear edge of species ranges, because these locations are ecologically more important, and because they should provide higher statistical power. Leading or rear range edges are the locations where populations are most sensitive to climate (Morley et al. 2017, Amburgey et al. 2018, Kleinhesselink and Adler 2018); accordingly, range edges also show the highest variability in population growth rates (Sexton et al. 2009, Csergő et al. 2017, Guyennon et al. 2023). From an ecological perspective, such high climate sensitivity provides an opportunity to understand range limit formation and to forecast range shifts (Parmesan 2006, Ehrlén and Morris 2015). From a statistical point of view, a higher sensitivity to climate increases statistical power.

We believe that the assumptions of our power analysis are likely to hold, as they reflect current ecological understanding. First, the assumption that populations experiencing the same average climate respond similarly to climate variation is well supported in the literature (Morley et al. 2017, Amburgey et al. 2018, Kleinhesselink and Adler 2018). Moreover, our power analysis is robust to relaxing this assumption (Fig. S6). Second, the assumption that residual variance (represented by *ε* in Eq. 2c) is constant across replicates also reflects current knowledge. Specifically, review (Sexton et al. 2009) and synthesis studies (Csergő et al. 2017, Guyennon et al. 2023) indicate that the variability of population growth rates should become larger only at range edges. Third, so long as the spatial replicates are sufficiently spaced (e.g., separated by dispersal), they should meet the assumption of statistical independence. In principle, statistical independence might not hold in response to unobserved large-scale drivers such as herbivores with large foraging ranges, or epidemic outbreaks. For this reason, we recommend testing for the presence of spatial autocorrelation in the residuals (see below).

To improve ecological inference, we suggest that researchers exploit the covariates that might mediate demographic responses to climate, such as soil conditions (Nicolè et al. 2011) or competition (Alexander et al. 2015). In single-species studies, these mediators cannot be taken into consideration, putting the external validity of the results into question. Spatial replication allows to quantify and test the importance of mediating factors. However, because the effect of these mediators is assessed by estimating an interaction, this requires a concomitant increase in sample size (Ch. 16: Gelman et al., 2020).

To detect spatial autocorrelations that might occur due to unexpected, large-scale processes such as epidemics, we suggest that spatial replicates should be sampled randomly rather than on a regular grid (Fortin et al. 1989). Detecting spatial autocorrelation is important, because it would indicate that the spatial replicates are not fully independent. A large spacing between groups of sites (e.g. 100 Km) would contribute to minimize such potential autocorrelation, while minimizing the climatic correlation among sites. This spacing might be possible for a distributed network of researchers, but it may be prohibitively expensive or logistically difficult for a single doctoral researcher. We suggest that, given the relatively small effect of climate correlation on statistical power, smaller spacing between populations might be a viable option in many real world situations.

Our simulations and analyses ignored the importance of microclimate which, however, might still be leveraged in spatially replicated studies. Recent studies highlight the importance of microtopography (Scherrer and Körner 2010), exposure (Ackerly et al. 2020), and vegetation type (Sanczuk et al. 2023) on plant responses to climate. It is unlikely that nearby sites experiencing different microclimates will also observe substantially different annual climatic anomalies. However, microclimates experience different absolute climatic conditions (e.g. minimum and maximum temperatures (Bennie et al. 2008). These conditions can be leveraged in studies that use physiological climatic predictors, such as growing degree days. While such physiological predictors require daily weather data, these can be measured via data loggers, or derived from gridded climatic data (e.g. (Maclean et al. 2019, Kearney and Porter 2020).The spatially replicated sampling we propose here is a practical solution to estimate climate-demography relationships and rapidly fill important knowledge gaps in the field of population ecology. However, monitoring 20 or more populations simultaneously is a large task for a single researcher. Such spatial replication might become more feasible through collaborative research networks (e.g. Villellas et al., 2021), or through unmanned aerial vehicles (UAV). UAV might make spatially replicated plant demographic data increasingly easy to obtain. For example, in both grassland and forest vegetation, UAV can identify individuals of a focal species (Schmidt et al. 2017, Bogdan et al. 2021, Allen et al. 2023), and quantify vegetation structure (Zhang et al. 2021, Coverdale and Davies 2023). We believe that the sampling choices we advocate in this article will contribute to the maturation of population ecology and its links to conservation science, functional ecology, and macro-ecology.

## Acknowledgements

We performed the analyses on the spatial correlation of climate using the High-Performance Computing Cluster EVE (a joint effort of both the Helmholtz Centre for Environmental Research - UFZ (http://www.ufz.de/) and the German Centre for Integrative Biodiversity Research (iDiv) Halle-Jena-Leipzig (http://www.idiv-biodiversity.de/).

## Data availability statement

The data and reproducible code for the analyses connected to this study are available at https://doi.org/10.5281/zenodo.8271041. We downloaded climatic data via CHELSA, dataset version 1.2, which is available at http://dx.doi.org/doi.10.5061/dryad.kd1d4.

## Ethics statement

We identified no ethical issues associated with this project.

## Author contributions

**Aldo Compagnoni**: Conceptualization (lead), data curation (lead), formal analysis (lead), investigation (lead), methodology (lead), resources (lead), software (lead), validation (lead), visualization (lead), writing – original draft (lead), writing – review & editing (equal). **Sanne Evers**: Conceptualization, data curation, formal analysis, investigation, methodology, resources, software, visualization, writing – review & editing (equal). **Tiffany M. Knight**: Conceptualization, funding acquisition (lead), methodology, project administration (lead), supervision (lead), writing – review & editing (equal).

