## Supplementary Material for "Spatial replication can best advance our understanding of population responses to climate"

### APPENDIX

#### Sensitivity of power estimates to variation in $\boldsymbol{\beta_1}$

To evaluate the sensitivity of power estimates to variation in the values of $\beta_1$, we modified the simulations presented in Eq. 2. We simulated a nested sampling design in which different populations are sampled in groups of three that experience identical $\beta_1$. In these simulations, equation 2a contains *ρ* values equal to one. In the first sensitivity analysis, each group of three populations has a different $\beta_1$ value, so that

$$\beta_{1_g} \sim Normal(\mu = 0.05, \sigma), \qquad \text{(S1)}$$

where $\beta_{1_g}$ are the effect sizes for group *g* of three populations, and the values suggest a mean $\beta_{1_g}$ value of 0.05 with a standard deviation of $\sigma$. We tested three values of $\sigma$: 0.007, 0.0125, and 0.025. Note that in the largest $\sigma$ value, over 5% of $\beta_{1_g}$ values are expected to be lower than zero. We explored the change in statistical power introduced by Eq. S1 by simulating three years of data, and a number of populations going from a minimum of three to a maximum of 48, in increments of three.

**Figure S1.** The correlation between temperature and precipitation anomalies decays slowly with distance. The left column shows how temperature (A) and precipitation (C) anomalies change with distance from a reference location located in Southern Florida. The red line and red shaded area show, respectively, the average fit and the 95% confidence interval of a polynomial local regression (loess) model relating the climatic correlation and the distance to the reference point. The right column shows heatmaps of correlations in temperature (B) and precipitation (D) anomalies between the reference location (red point) and the remaining points for which we sampled climatic data.

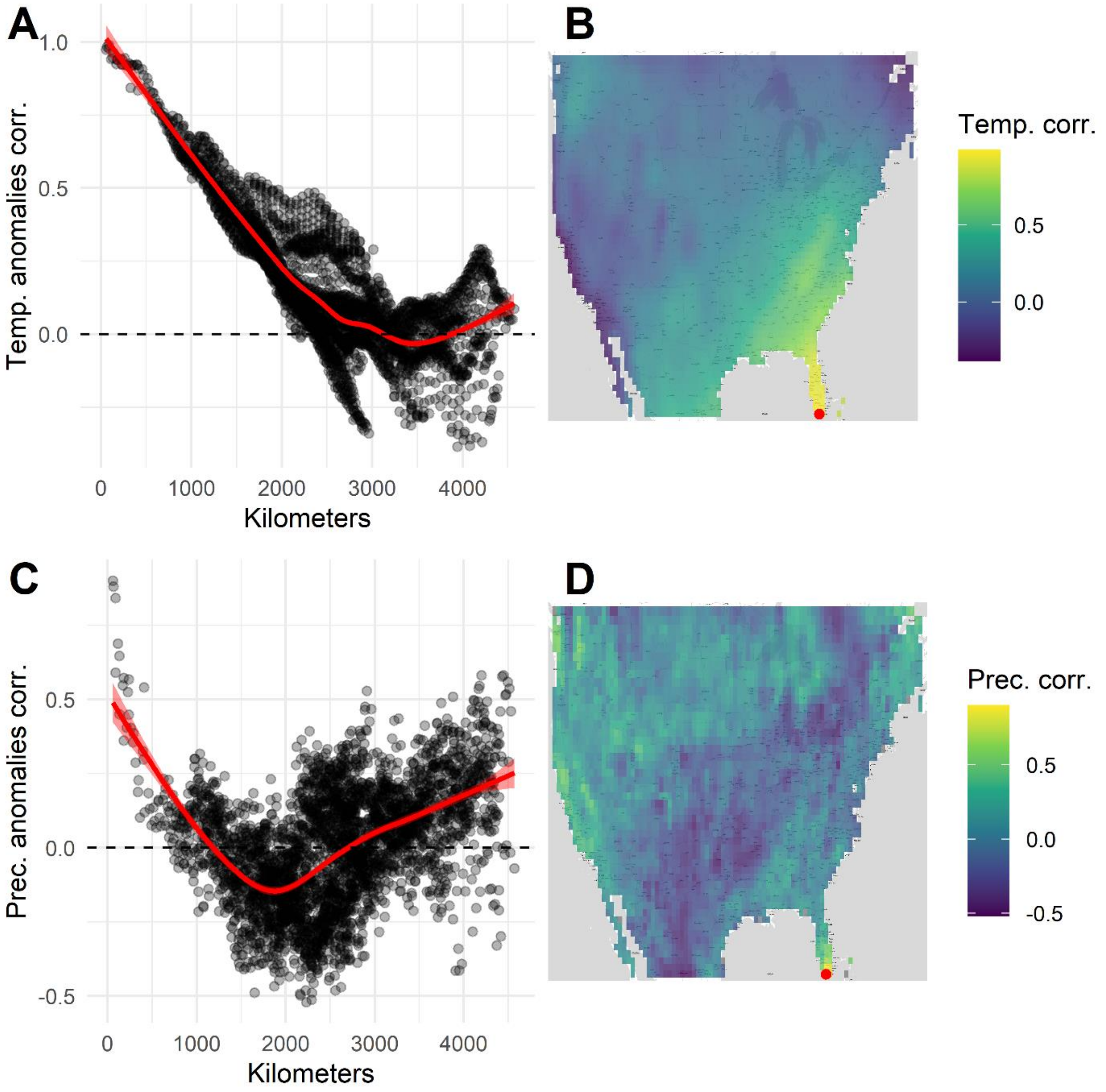

**Figure S2.** The correlation between temperature and precipitation anomalies decays slowly with distance. The left column shows how temperature (A) and precipitation (C) anomalies change with distance from a reference location located in coastal California. The red line and red shaded area show, respectively, the average fit and the 95% confidence interval of a polynomial local regression (loess) model relating the climatic correlation and the distance to the reference point. The right column shows heatmaps of correlations in temperature (B) and precipitation (D) anomalies between the reference location (red point) and the remaining points for which we sampled climatic data.

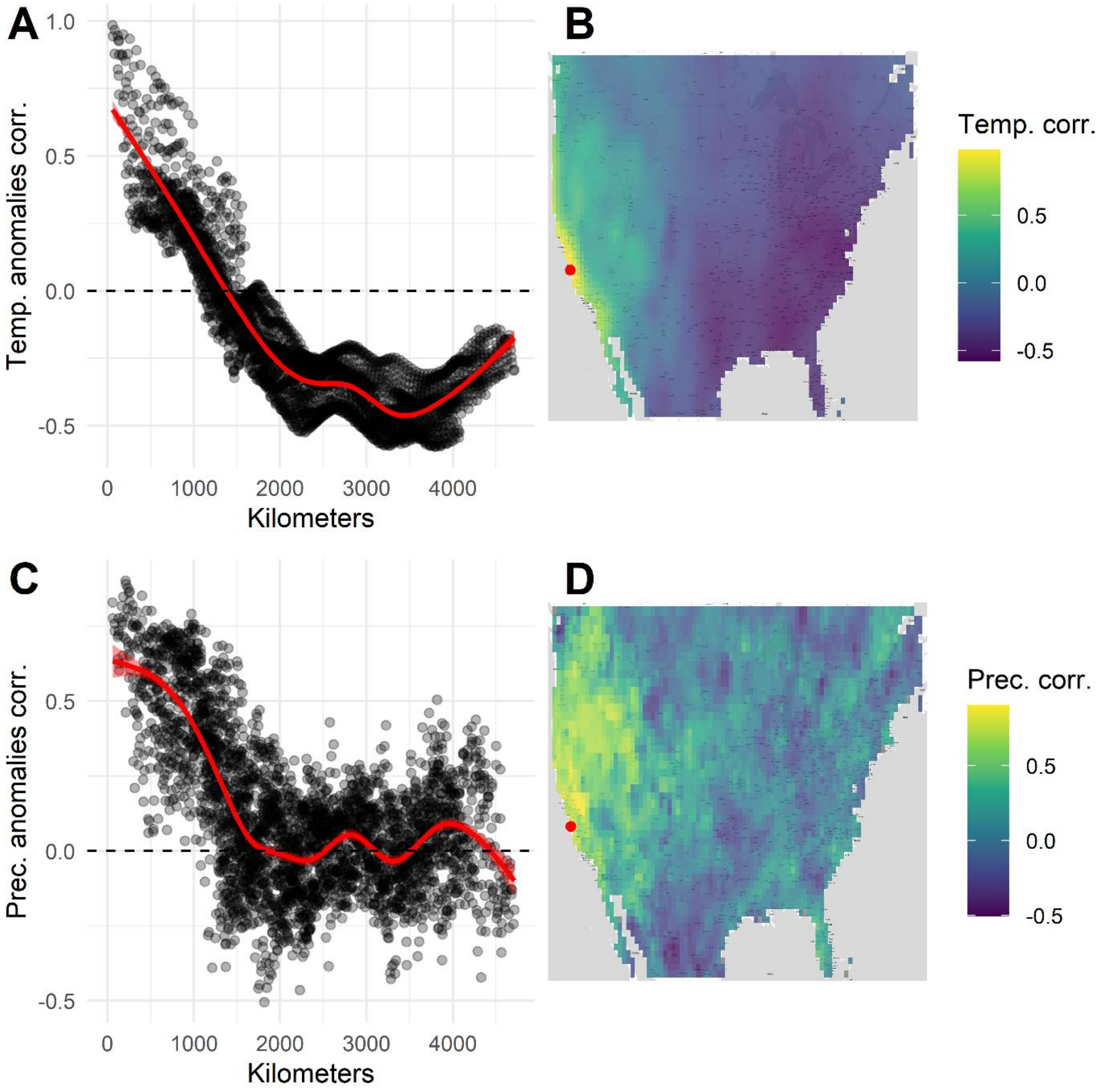

**Figure S3.** The correlation between temperature and precipitation anomalies decays slowly with distance. The left column shows how temperature (A) and precipitation (C) anomalies change with distance from a reference location located near the city of Boston. The red line and red shaded area show, respectively, the average fit and the 95% confidence interval of a polynomial local regression (loess) model relating the climatic correlation and the distance to the reference point. The right column shows heatmaps of correlations in temperature (B) and precipitation (D) anomalies between the reference location (red point) and the remaining points for which we sampled climatic data.

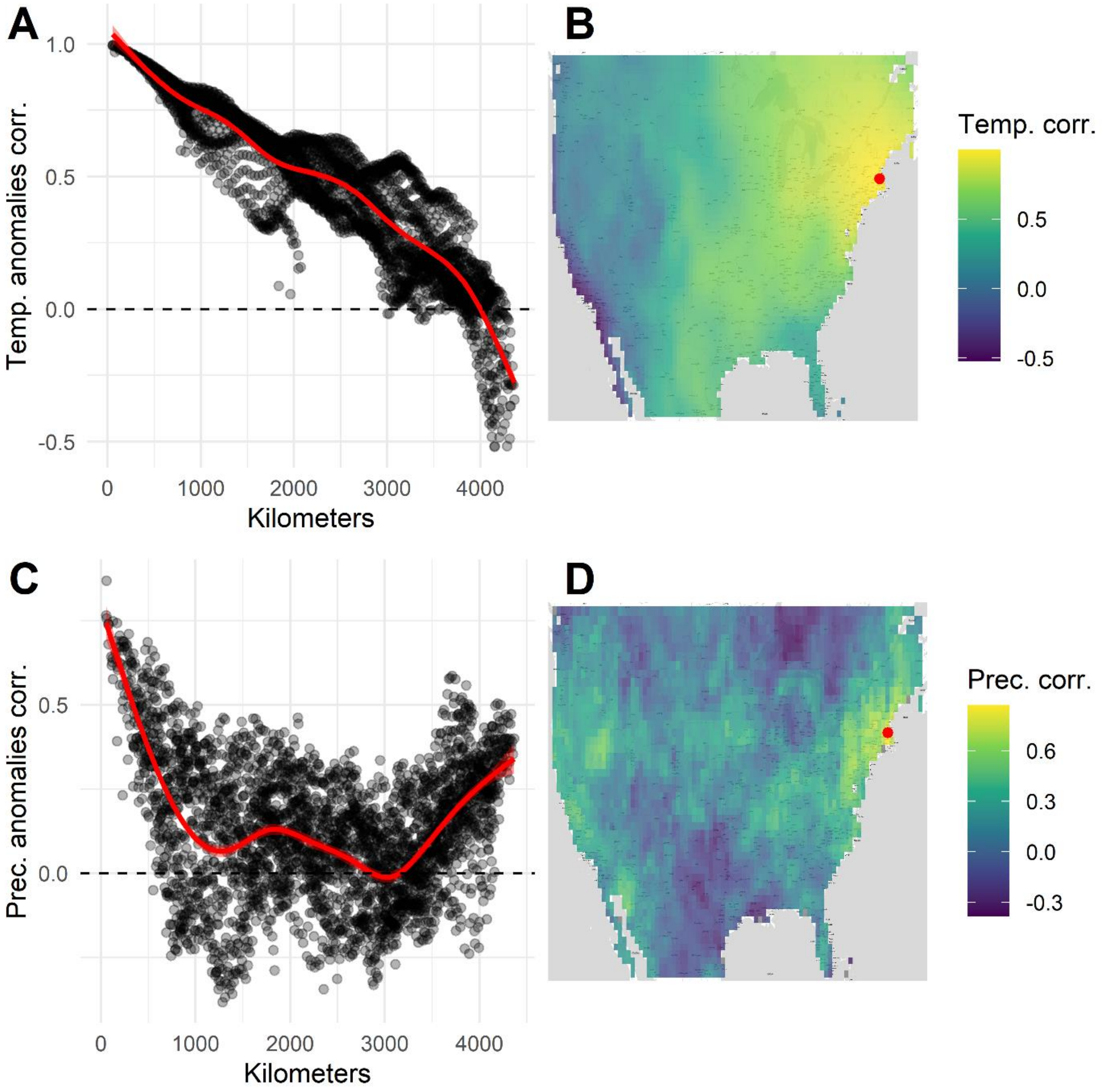

**Figure S4.** The correlation between temperature and precipitation anomalies decays slowly with distance. The left column shows how temperature (A) and precipitation (C) anomalies change with distance from a reference location located near the city of Seattle. The red line and red shaded area show, respectively, the average fit and the 95% confidence interval of a polynomial local regression (loess) model relating the climatic correlation and the distance to the reference point. The right column shows heatmaps of correlations in temperature (B) and precipitation (D) anomalies between the reference location (red point) and the remaining points for which we sampled climatic data.

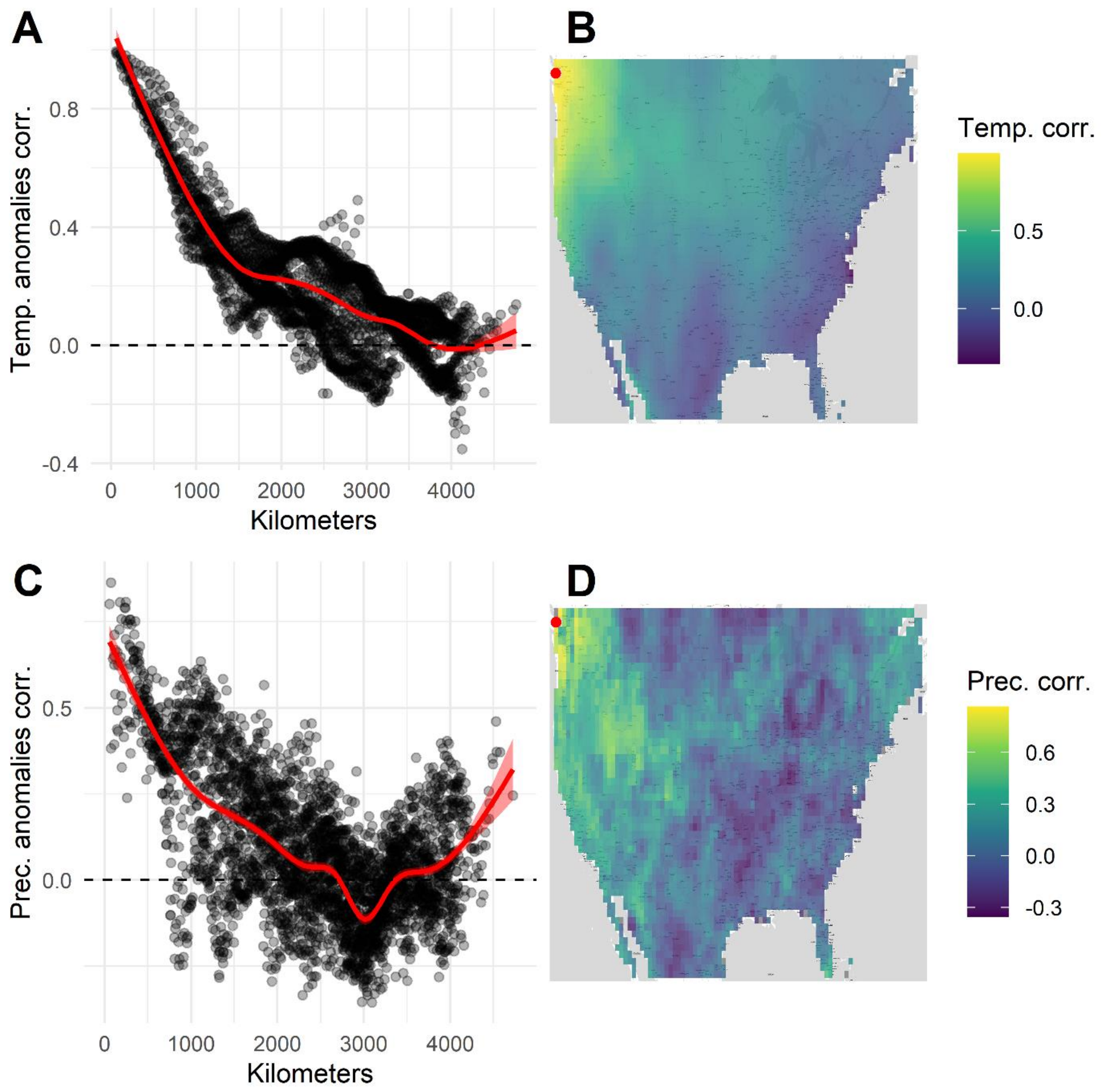

**Figure S5.** The correlation between temperature and precipitation anomalies decays slowly with distance. The left column shows how temperature (A) and precipitation (C) anomalies change with distance from a reference location located in the midwest. The red line and red shaded area show, respectively, the average fit and the 95% confidence interval of a polynomial local regression (loess) model relating the climatic correlation and the distance to the reference point. The right column shows heatmaps of correlations in temperature (B) and precipitation (D) anomalies between the reference location (red point) and the remaining points for which we sampled climatic data.

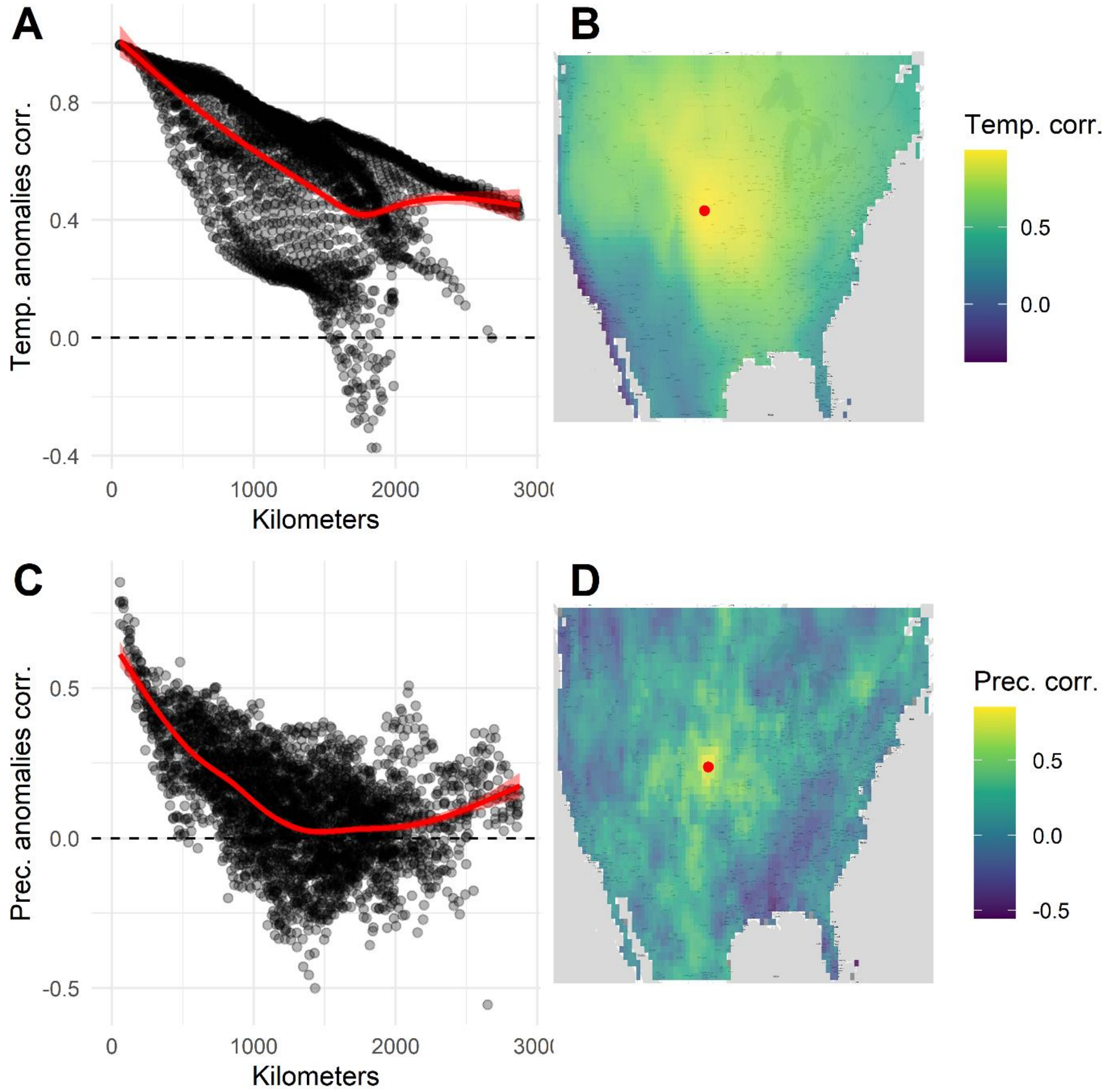

**Figure S6.** The lower temporal correlation between two climatic anomalies allows to sample a larger absolute range of anomalies. Bivariate plot representing the range of climate anomalies sampled (y-axis) at two hypothetical sites, as a function of years sampled (x-axis). The color of dots shows the correlation of climate anomalies at these two hypothetical sites. A correlation of one implies that the two sites experience identical climatic anomalies each year.

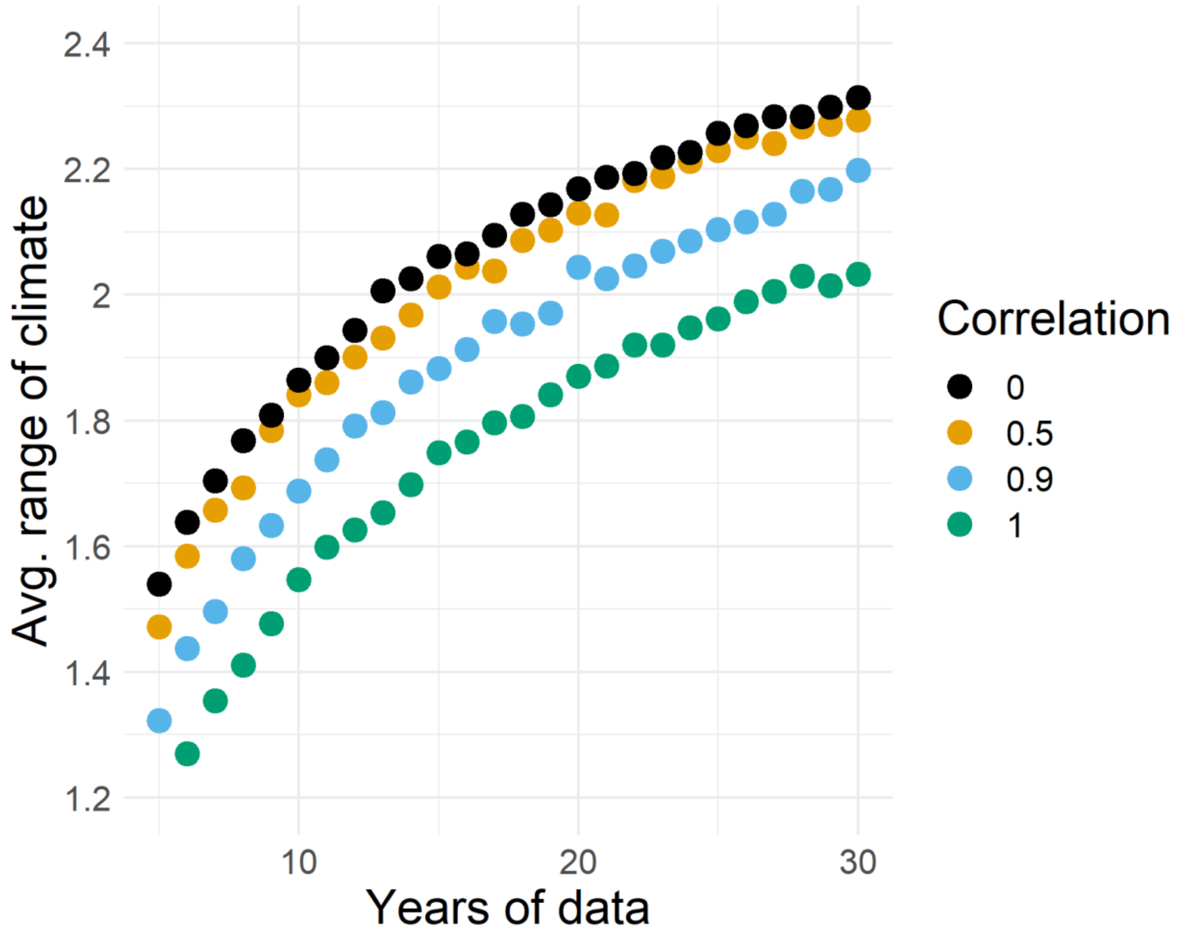

**Figure S7.** Variance in the effect of climate among datasets does not substantially impact statistical power. Plot showing statistical power (y-axis) as a function of spatial replication (x-axis). Colors show the standard deviation in the effect of climate, $\beta_1$, which in these simulations was a random variable drawn from a normal distribution.

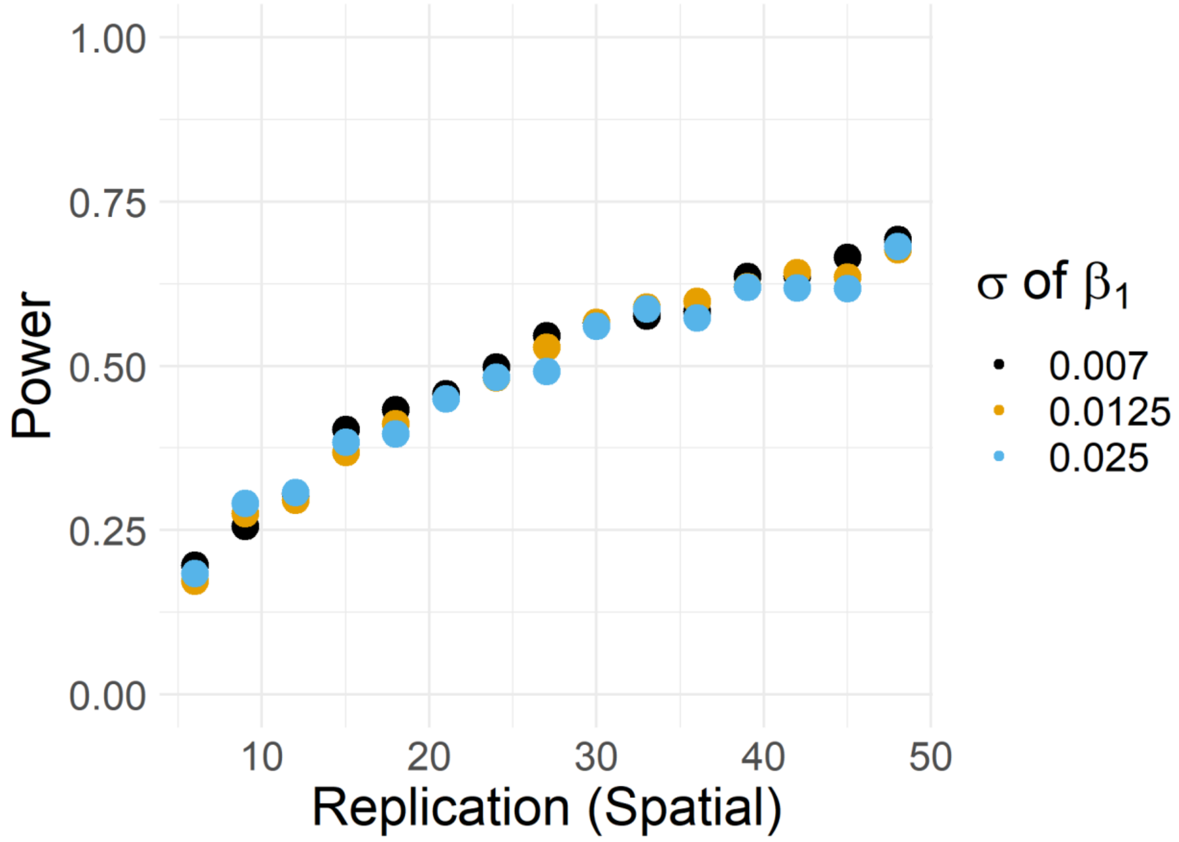
